# Predation and the Evolution of Island Bird Plumage Colouration: Experimental Insights from Island and Mainland Environments

**DOI:** 10.64898/2026.01.31.703000

**Authors:** Ana V. Leitão, Cristina D. Alonso Moya, Ricardo J. Lopes, Raquel Ponti, Rita Covas, Claire Doutrelant

## Abstract

Islands serve as natural laboratories for exploring evolutionary processes, often fostering unique species through their isolation and distinct ecological conditions. These environments present opportunities to study how a range of selective pressures shape biodiversity. Bird plumage colouration is one trait that has shown to consistently change in island populations, and predation has been hypothesized to influence these differences. While animals often face a trade-off between signalling to conspecifics and avoiding detection by predators, the role of predation in shaping conspicuousness remains underexplored experimentally. In this study, we asked how predation pressure differs between insular and mainland habitats, and whether predation risk covaries with conspicuousness of male and female birds across environments. In a field experiment, we investigated predation rates using 3D-printed models painted to represent both sexes of 12 bird species from three archipelagos (Madeira, Azores, and Canary Islands) and their closest mainland relatives. These models were deployed in the species’ natural environments to measure hit rates (a proxy for predation risk), accounting for factors that influence prey detectability, such as colour of the models, background contrast, and vegetation. We found that models on the islands experienced less hits compared to those on the mainland, while sexual dichromatic models were more likely to be dislodged on the mainland. In addition, for mainland sites, increased chromatic contrast correlated with a higher probability of dislodgment, suggesting that more conspicuous models were more likely to be hit. These results highlight that while predation constrains conspicuousness, other ecological and evolutionary factors likely drive the reduced plumage colouration observed in island birds. Our research offers experimental insights into how predation interacts with conspicuous traits in shaping plumage colouration in birds.

## Introduction

Oceanic islands offer a unique window into the processes of evolution and adaptation, due to their simplified ecological settings, small and genetically limited populations and reduced species diversity (MacArthur and Wilson 1963, Kier, Kreft et al. 2009, Whittaker, Fernandez-Palacios et al. 2017). These conditions promote the evolution of distinctive traits among insular species, often differing markedly from their continental relatives in morphology (e.g., dwarfism or gigantism), life history (e.g., lower fecundity), and behaviour (e.g., reduced wariness) (Lomolino 2005, Covas 2012, Novosolov, Raia et al. 2013, Cooper Jr, Pyron et al. 2014).

Among these traits, visual signals such as plumage colouration, play a central role in intraspecific communication and species recognition, making them particularly relevant for understanding evolutionary dynamics in isolated populations, such as island environments. Yet, the effects of insularity on key traits like animal signalling remain relatively underexplored (Jezierski, Smith et al. 2023). Some studies reported reduced ornamentation in island species (Grant 1965, Fitzpatrick 1998, Figuerola and Green 2000, Roulin and Salamin 2010, Doutrelant, Paquet et al. 2016), while others have linked reduced predation to increased colouration (Runemark, Brydegaard et al. 2014, Bliard, Paquet et al. 2020). These findings indicate that island birds may be duller than mainland species, but the role of predation in shaping these patterns remains unresolved. Conspicuous colouration, while beneficial for communication, can increase detectability to visually oriented predators such as raptors, potentially constraining the evolution of elaborate colours (Huhta, Rytkönen et al. 2003, Husak, Macedonia et al. 2006, Vignieri, Larson et al. 2010).

Predation is a key ecological force shaping trait evolution, and islands are often assumed to experience reduced predation pressure due to lower predator richness (MacArthur and Wilson 1967, Beauchamp 2004). In some cases, islands lack predators entirely, which clearly supports this assumption. However, most islands retain some predators, and it remains unclear whether less species of predators translates into lower overall predator abundance or reduced predation risk. According to the island syndrome hypothesis, lower predator diversity could broaden predator niches, potentially maintaining or even increasing predation pressure (Carlquist 1974, McNab 1994). Much of the perceived reduction in predation pressure on islands is inferred from comparisons of behavioural or ecological traits between island and mainland populations, such as weaker anti-predator behaviours and greater naivety in island species (Blumstein and Daniel 2005, Cooper Jr, Pyron et al. 2014, McWaters and Pangle 2021), while direct experimental evidence testing whether predation pressure is consistently reduced on islands is still lacking.

In sexually dichromatic species, males typically display more elaborate colouration than females, due to differing selective pressures, with stronger sexual selection acting on males, and greater natural selection acting on females (Andersson 1994). As a result, males are generally expected to incur a higher predation risk than females because of their greater conspicuousness (Dale, Dey et al. 2015, Delhey, Valcu et al. 2023). However, some studies have found increased attacks on more cryptic individuals instead (Götmark 1993, Götmark and Unger 1994, Ruiz-Rodríguez, Avilés et al. 2013, Cain, Hall et al. 2019), suggesting that the relationship between colouration and predation risk is highly context-dependent. Importantly, in monochromatic species, males and females are similarly coloured, whereas in dichromatic species, conspicuousness is unevenly distributed between sexes, a heterogeneity in visual signals that may influence the probability of being encountered by predators and the overall predation risk, independent of whether predators recognise species identity.

In this study, we asked whether predation pressure differs between mainland and islands and whether this variation is associated with differences in bird colouration. We used an experimental approach with 3D-printed bird models of adult males and females from 12 species (6 island-mainland pairs) across the Atlantic Islands of Macaronesia and mainland Portugal. We quantified predation risk using the hit rate, defined as the proportion of models that were dislodged during deployment. We deployed these 3D models, representing island-mainland species, in their natural habitats. The use of model animals (clay or 3D-printed) has proven effective in assessing predator responses to prey traits in natural settings (Bateman, Fleming et al. 2017, Cain, Hall et al. 2019, Whiting, Holland et al. 2022), as it allows to isolate specific visual cues, such as colouration or size, and systematically test their influence on predation risk, while controlling for behaviour and movement. Building on established experimental and visual-modelling approaches using model prey (Stuart-Fox, Moussalli et al. 2003, Cain, Hall et al. 2019, Whiting, Holland et al. 2022), a particularly novel aspect of our approach is the application of these methods with multiple species across different islands and mainland locations to experimentally assess long-standing assumption that island species experience reduced predation. Overall, we expected to find that (a) hit rates would be higher on the mainland than on islands, (b) more colourful and contrasting models would elicit higher hit rates than duller ones, and (c) male models would experience more hits than female models, reflecting higher conspicuousness.

## Methods

### Study Species

The study involved 12 passerine species, arranged in 6 pairs of island endemics and their closest mainland relatives. At the time of the study, we selected only taxa recognised as full species, although some endemics have since been elevated from subspecies status (e.g. *Fringilla canariensis* and *Fringilla madeirensis*). The different species differ in the levels of habitat and sexual dimorphism (Supplementary Table S1). Three species are monomorphic (*Phylloscopus canariensis, Phylloscopus ibericus, Pyrrhula murina*), and all the others are dimorphic with variation in the degree of sexual differences.

The island endemics studied inhabit the Atlantic Islands of Macaronesia, specifically the archipelagos of Madeira (Madeira), Azores (São Miguel), and the Canary Islands (Tenerife and Fuerteventura). This region exhibits a wide range of biogeographical characteristics, including island size (293 km2 – 2034km2), varying distance from the mainland (ranging from less than 100 km to 1,365 km), latitude, geological age (0.25-26 million years), and altitude (130-3700 m above sea level). These factors contribute to a wide array of ecological conditions, including strong climatic gradients and varying predator assemblages: from a single diurnal bird of prey in the Azores, to 4 in Madeira and up to 5 in the Canary Islands, with numbers varying among islands within each archipelago. Such variation fosters rich habitat diversity and supports a significant level of endemism, with approximately 25% of species endemic to the region (Illera, Rando et al. 2012, Florencio, Patiño et al. 2021). The corresponding mainland species have wide distribution ranges (Supplementary Table S1), and for comparison, we selected populations from mainland Portugal, which occur at latitudes as similar as possible to those of the island endemics.

### Spectrophotometry

Reflectance measurements were taken from (1) museum bird specimens from the target species to assess the degree of colour similarity between models and plumage, (2) painted 3D models and (3) background measurements from natural vegetation and structures surrounding the models, to analyse colour contrast.

Reflectance spectra from bird specimens were measured from three male and three female adult specimens, except where fewer specimens were available. The specimens were sourced from the British Natural History Museum, the American Natural History Museum, the Naturalis Museum of Leiden in the Netherlands, and Museum of Natural History and Science of the University of Porto, Portugal. Measurements were taken in the bird-visible wavelength range (300-700 nm), using either a USB 4000 spectrometer with a PX2 Xenon light source or a Jaz spectrophotometer (Ocean Optics, Largo, FL) and a 45-degree angle probe (UV-VIS fibre-optic reflectance). The probe end excluded all ambient light and maintained a 2mm fixed distance from the surface. Reflectance was calibrated relative to a dark and white standard (WS-2 reference tile, Avantes). Measurements were made with the reflectance probe gently touching perpendicularly the feathers, the surface of the acrylic model or the vegetation. For each species and sex, we took reflectance measurements from visually prominent body regions, following a consistent approach similar to Doutrelant et al. (2016). We measured colouration in at least eight body regions: head, breast, belly, mantle, uropygial area, shoulder, wing, tail, and undertail. For species with additional distinct patches, these were also measured. We took five measurements for each colour patch on bird specimens and 3D models, and three measurements for each type of vegetation or structure (see below).

### Model construction

The use of 3D-printed prey models to quantify predation risk follows approaches developed and validated in previous studies (Cain, Hall et al. 2019, Walker and Humphries 2019). Models of the 12 target species were created using 3D printing, based on digitisations produced through 3D photogrammetry of taxidermic specimens, and painted with colours that replicate the natural plumage colours for each species and sex, also based on specimens from museums (Figure 1).

**Figure 1.**
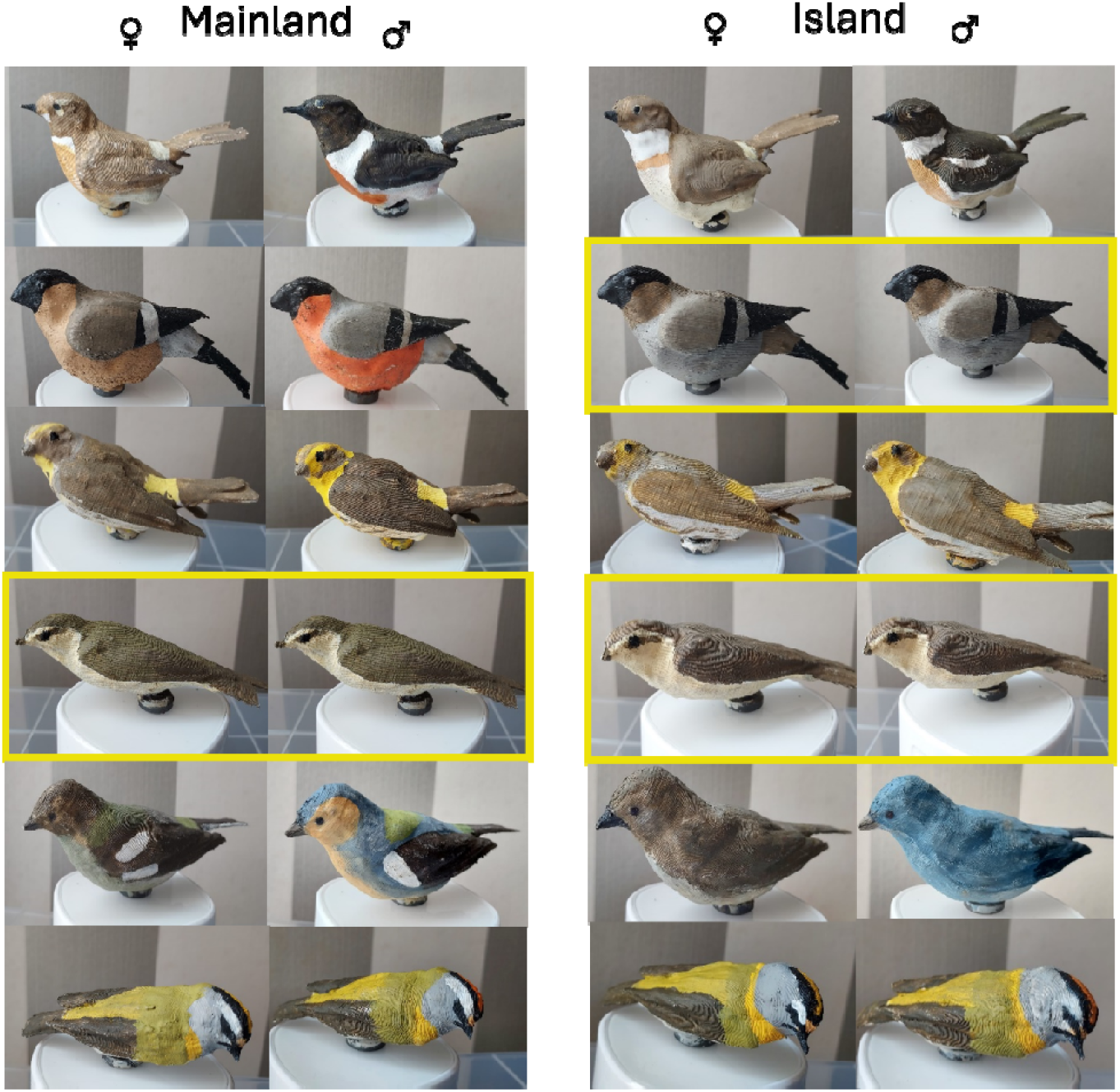
3D-printed and hand painted models of the target species. On the left column the mainland and on the right the island endemic species. Within each column the left panels depict the females and the right ones the males. For monochromatic species, the same photograph is repeated and marked with a yellow square. From top to bottom, left to right: European stonechat (*Saxicola rubicola*), Fuerteventura stonechat (*Saxicola dacotiae*), Eurasian bullfinch (*Pyrrhula pyrrhula*), Azores bullfinch (*Pyrrhula murina*), European serin (*Serinus serinus*), Atlantic Canary (*Serinus canaria)*, Iberian Chiffchaff (*Phylloscopus ibericus*), Canary Islands Chiffchaff (*Phylloscopus canariensis)*, Common chaffinch (*Fringilla coelebs)*, Blue Chaffinch (*Fringilla teydea)*, Common firecrest (*Regulus ignicapilla)*, Madeira firecrest (*Regulus madeirensis)*. More information on each species and their distribution can be found on Supplementary Table S1.

The taxidermic specimens used for the photogrammetry were housed at the Museum of Natural History and Science of the University of Porto, Portugal. The hardware setup for capturing 360-degree images of the specimens was adapted from Medina et al. (2020) (Supplementary Figure S2). A Canon EF-M M5 APS-C camera with a lens Canon EF-M 32mm F1.4 STM (Canon Inc., Japan), mounted on a Manfrotto 190 tripod (Manfrotto, Italy) and controlled remotely was positioned inside a Foldio3 (Orangemonkie Inc., California) foldable photo studio with a matte white background. Each specimen was placed on a transparent stand on top of a turntable, inside the photo studio. Consistent lighting (5600 K) was ensured by the Foldio3 triple LED strips and two external LED light sources positioned symmetrically on either side of the setup. For each imaging session, the camera was configured with an f/11 aperture, ISO 400, manual focus, and a continuous slow-shooting mode. Photographs were taken continuously using the remote control while the turntable rotated at a constant speed, capturing approximately 100 images for a full 360-degree rotation. This process was repeated three times: at the specimen’s central level, 45 degrees below, and 45 degrees above (Supplementary Figure S2).

Post-processing of the images involved adjusting colours and image dimension and removing background using Adobe Lightroom and Adobe Photoshop (Adobe Inc., California, EUA). The 3D models were constructed using Metashape (Agisoft LLC, St. Petersburg, RU) by aligning the 360-degree images, generating a dense cloud, creating a mesh, and adding texture. Final mesh refinements and corrections were performed in Blender (Blender Foundation, Amsterdam, NL). Once finalized, the models were commercially printed in white polylactic acid (PLA) by 3D Filamentos (Matosinhos, Portugal). Due to similarities in body shape within species pairs, approximately 45 plain models per genus were printed. The printed models were then hand-painted to replicate the natural plumage of the target species. A colour palette was generated for each sex and species by extracting tones from high-resolution photographs. Acrylic paints were then selected, mixed, and refined using spectrophotometry to closely match the reflectance spectra of museum specimens, accounting for UV wavelengths, which were assessed with a spectrophotometer and were compared with the specimen colours through avian visual models (see below). Models were painted to represent each species and each sex (where applicable, for dichromatic species). For sexually monochromatic species, such as *Pyrrhula murina* (Ramos 1997), and the *Phylloscopus* species (Gordo, Arroyo et al. 2016), we assumed no distinction between sexes. The models were categorized into “Model types” based on their origin and sex: female island, male island, female mainland, male mainland; for monochromatic species, models were categorised as either mainland or island species (Figure 1).

### Visual Modelling

We analysed the colours using avian visual models applied to spectral reflectance data for comparison between patches of 3D models and their museum specimen counterparts and model detectability by visual predators in their natural environment, by calculating chromatic and achromatic contrast between each model type and the natural background where they were placed during the experiment.

All reflectance data was visualised, checked for quality, adjusted for minor negative values, and smoothed using smoothing at 0.2 with the plotsmooth function in the *pavo* package (version 2.8.0, Maia, Gruson et al. 2019) in R (version 4.3.0, R Core Team 2024).

Colour reflection was modelled from the perspective of a diurnal predator raptor, using the receptor-noise limited (RNL) model of Vorobyev et al. (1998) and Vorobyev et al. (2001), to estimate the perceptual distance between coloured stimuli (colour discrimination). This model assumes that colour discrimination is determined by the noise in the photoreceptors of a visual system, the relative abundance of these photoreceptors and the opponent mechanisms by which these photoreceptor signals are processed.

First, we estimated photoreceptor quantum catches, assuming independent process of the chromatic and the achromatic component of the colour. We quantified the stimulation of receptors using the function *vismodel* in *pavo* package. Birds possess tetrachromatic vision, with four single-cone types. Because the visual systems of most avian predators, including raptors, are violet-sensitive (V-type) (Ödeen and Håstad 2013), we employed an average V-type visual system with peak sensitivities from Endler and Mielke (2005): VS= 412; S= 452, M= 505, L= 565. Quantum catch values for each photoreceptor were computed under standard daylight (D65) illumination, incorporating von Kries chromatic adaptation and the ideal background reflectance.

To estimate the contrast between each model type and the natural background on which they were placed during the experiment, we calculated the chromatic and achromatic colour distances between model colour patches and site-specific background reflectance using the visual system described above. Colour contrasts are expressed in units of JNDs (just noticeable differences), where 1 JND is the theoretical threshold of colour discrimination. These distances indicate to what extent the different patches resemble natural backgrounds in colour, with higher values indicating stronger contrast.

Background colours were taken in the field and grouped in two categories: green (leaves, grass, bushes), and brown (leaves, soil, and bark) (see reflectance spectra in Supplementary Figure S1). Chromatic and achromatic contrasts were then derived using the *coldist* function in *pavo*. The noise-to-signal ratio of each cone type was calculated based on formula 10 in Vorobyev et al. (1998) from the average cone proportions of V-type birds from Hart (2001: VS = 0.4, SWS = 0.75, MWS = 1.2, LWS =1), and a Weber fraction of 0.1 for the L cone (Olsson, Lind et al. 2018). Achromatic contrast was derived from the double-cone, which mediates luminance perception in birds (Hart 2001). Double-cone quantum catches were modelled using the double-cone sensitivity function for the Blue tit *Cyanistes caeruleus* implemented in *vismodel* function.

For each model species type we calculated the chromatic and achromatic contrast values for all measured colour patches. For subsequent analysis, we used only the patch with the highest contrast. This provided a single measure of overall conspicuousness for each model species, representing the maximum contrast over the full range of natural backgrounds (green and brown).

We also calculated the chromatic contrast between the bird specimen plumage and the painted models against natural background, using a similar approach described above. We confirmed a strong correspondence in chromatic contrast between the 3D models and the bird specimen plumage (Supplementary Figure S3).

### Experimental design

The experimental study took place in the locations where the island or the mainland species occurred, i.e. in the Atlantic Islands of Macaronesia, as well as mainland Portugal (Figure 2). The experiments were conducted from March 10 to June 27, 2023 (during pre-breeding of the species, see table S1). Pilot fieldwork was carried in 2022, in the Azores and mainland Portugal, involving only one pair-species (*Pyrrhula*), and followed a different experimental design, which was later revised to the standardized protocol described below. As such, the 2022 data were not incorporated in the visual modelling or statistical analyses, and the final dataset and results derive exclusively from the 2023 experiments.

**Figure 2.**
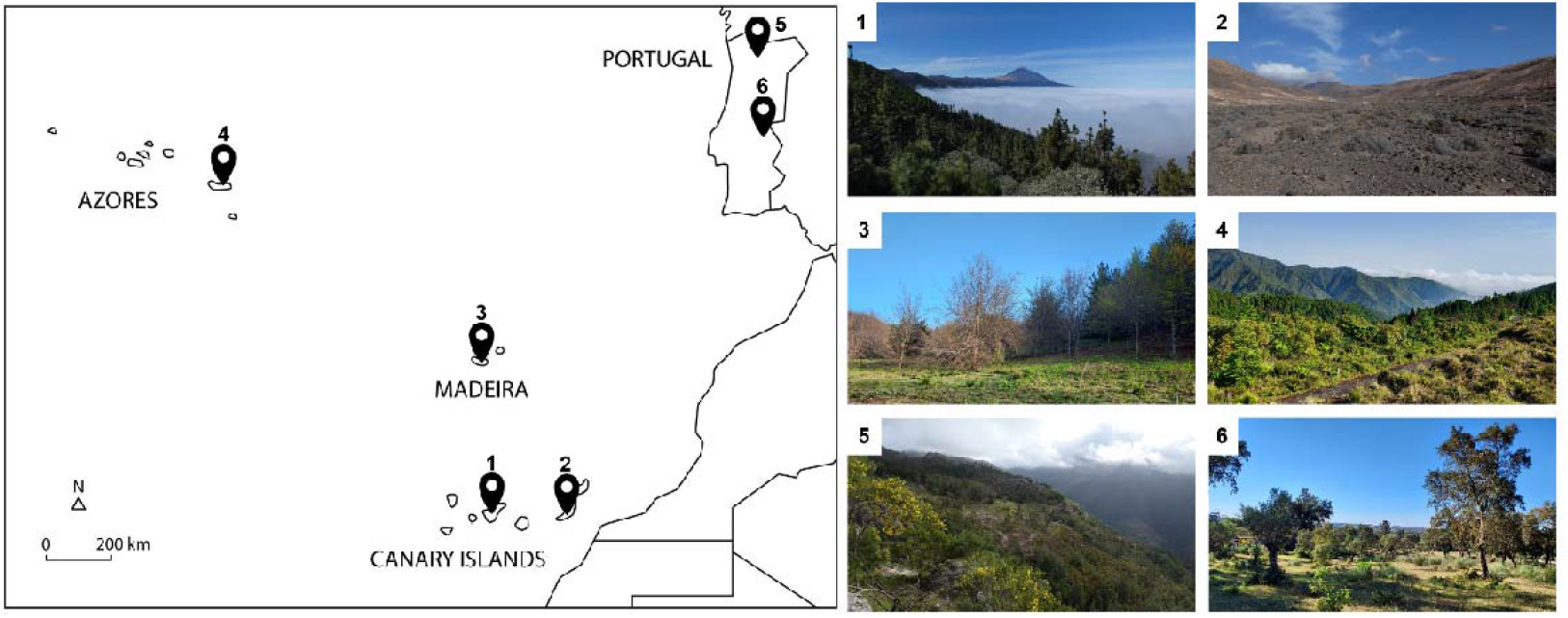
Illustration map of field study sites where the study species also occur (left) and photos of the different locations (right). Islands: Tenerife (1), and Fuerteventura (2; Canary Islands); Madeira (3; Madeira), São Miguel (4; Azores); Mainland: Campo do Gerês (5) and Portalegre (6; Portugal).

Transect locations were selected in areas where the target species was commonly observed, typically in short herbaceous vegetation, low shrub patches, or open ground within forested or semi-open habitats. These microhabitats resembled the substrates where the birds commonly forage and ensured that models were visible and accessible to potential predators. The precise GPS coordinates for each transect are provided in the dataset.

This experimental layout follows established model-prey deployment designs used to quantify predator attack rates in the field (Stuart-Fox, Moussalli et al. 2003, Cain, Hall et al. 2019). For each pair-species and at each location (island/mainland), we set 3 to 6 transects (Supplementary Table S2), each containing approximately 40 models. For monochromatic species (e.g. *Phylloscopus* sp.), transects included 20 island model species and 20 mainland model species (N=40). For dichromatic species, transects included 10 female and 10 male models for both mainland and island species (N=40). For the specific case of the *Pyrrhula*, we set 30 models, 10 for the monochromatic island species and 10 for each sex of the mainland species. The number of transects varied across sites and species due to logistical constraints.

Each model was marked with a unique number on the ventral side and was fitted with a magnet and a fishing line affixed to its ventral surface. This allowed secure attachment to a metal nail that was placed on the ground, preventing wind displacement and reducing the likelihood of long-distance movement by predators, while still enabling potential predators to interact with and dislodge the model from the magnet.

Transects were set up early in the morning, with models placed at 2-m intervals on short vegetation, to simulate foraging birds (Figure 3). For each transect we recorded the time of setup, coordinates, weather conditions, and the number of each model. Models were placed in alternating order (female, male, island, mainland, Figure 3), and were checked every 24h for signs of predator attack, which consisted of being dislodged from the nail, i.e. being hit. Each transect was left for 48h and then moved to another site more than 100 m away. To minimise disturbance, we did not visit transects between placement, checking and collection.

**Figure 3.**
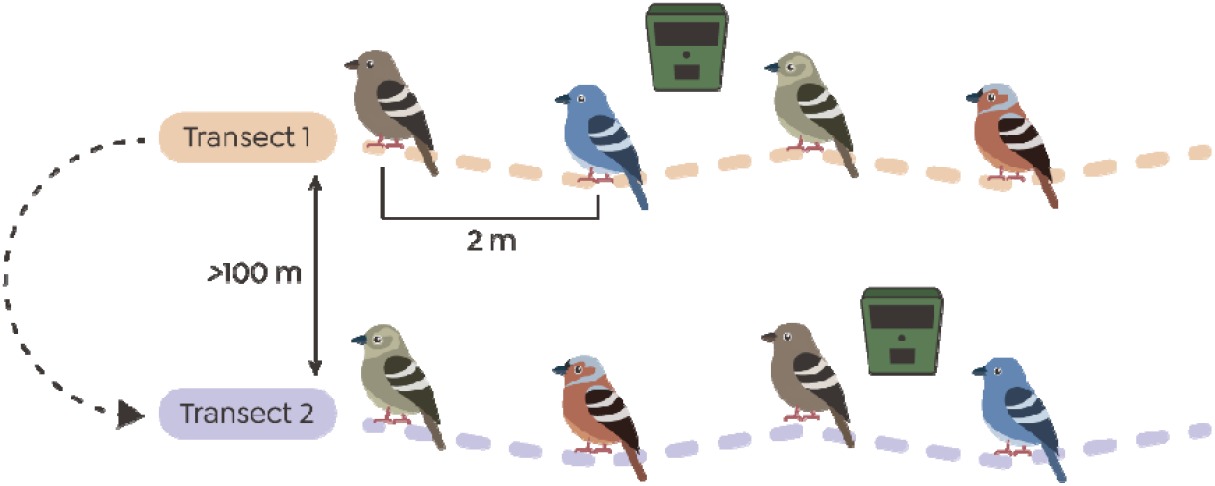
Experimental setup. In this example we show a dichromatic species pair: *Fringilla coelebs* (mainland chaffinch) and *Fringilla teydea* (island chaffinch). Model birds were positioned 2 meters apart in the following sequence: mainland female, mainland female, island female, and island male. A subset of models was monitored using camera traps to identify potential predators. Each transect was relocated every 48 hours to a new site at least 100 meters from the previous location.

To identify putative predators, we used 19 camera traps (14 cameras Browning Command Ops Elite 20, 3 cameras Bushnell, 1 camera Victure HC300, 1 Moultrie A900) triggered by an infrared motion-and-heat detector to obtain a sample of observational data on predator activity near the models during each field experiment. The cameras were positioned more than 1m away from the models and were active 24 hours a day. They were set to the highest sensitivity and automatically took photo or video when the infrared sensors detected motion, with most of the cameras having a 0.3s trigger speed (Browning Command Ops Elite 20). A small portion of the cameras failed to trigger when the models were hit (see results).

In total we performed 40 transect days on the mainland, and 42 transect days on the islands. In four transect-days we observed on the camera traps a high number of false hits due to local cattle, so we excluded the full transects from the analysis, resulting in 38 transect days in the islands and 40 transect days on the mainland (Supplementary Table S2). In 4 individual cases we detected that models were hit by non-predators (cattle and rabbits) or during the night (which presumably were not affected by model colour), and for that reason were excluded.

### Model background contrast and vegetation cover

To characterise the background environment, we measured the spectral reflectance of surrounding vegetation at all locations where experiments were conducted. Measurement of background spectral reflectance followed the same protocol as that used for plumage and model reflectance. Backgrounds were sampled across multiple habitat types, including forest understory, shrubland, and open ground areas, on both island and mainland sites. Within each habitat, we selected prominent leaves, bushes, moss, rocks, and soil patches surrounding the 3D models. We collected a total of 123 background measurements from 41 structures, with three measurements per structure. These data were combined by averaging the reflectance values of the most predominant colours, green and brown (Supplementary Figure S1). Chromatic contrasts between green and brown backgrounds were strongly correlated (r = 0.85), as were achromatic contrasts between green and brown backgrounds (r = 0.96). Visual modelling revealed that the highest chromatic contrast was generally against green backgrounds, followed by brown backgrounds (Supplementary Table S3). Overall, male models, particularly the *Fringilla, Regulus* and *Pyrrhula*, had the highest chromatic contrast between the most colourful patch and the background. In contrast, models of *Phylloscopus* and *Serinus* displayed the lowest levels of chromatic contrast (Supplementary Table S3).

To characterise the vegetation cover, we extracted Normalised Difference Vegetation Index (NDVI) from satellite-based raster files. NDVI quantifies vegetation greenness and estimates vegetation density, and it is calculated from spectrometric data. NDVI values range from - 0.08 to +0.92, where low values (≤0.1) indicate barren areas such as rock or sand; moderate values (0.2–0.5) correspond to sparse vegetation, such as grasslands or senescing crops; and high values (0.6–0.9) represent dense vegetation, typical of forests or crops at peak growth. NDVI data was retrieved from Copernicus Land Monitoring Service (Copernicus 2021). We obtained the data from a period of 10 days (version 2), at 300 m resolution. For each transect, we used the NDVI layer corresponding to the date of the experiment sampling, covering 12 time points from March to June 2023.

### Statistical analysis

In this study, each model check conducted every 24 hours was treated as a separate data point, with the outcome recorded as either “yes” or “no” depending on whether the model was hit. We first built a simple model (model 1) testing for an insularity effect on dislodgment rate (as a proxy for attack rates). Having found an effect (see results), we built additional models (model 2a, 2b, and model 3) for additional influence colour contrast and sex of the model (see below for more details). In all these models we fitted a generalized linear mixed-effects model with a binomial family and “Dislodged” (yes/no) as the response variable and included as a fixed effect “Transect day” (1^st^ or 2^nd^ day) to account for potential day effects, and the random effects “Transect number” to account for spatial variation and repeated measures within the same site, and “Pair” (the pair number of the island endemics and their closest mainland relative) to account for the phylogenetic paired structure of the analyses.

For model 1, we tested whether dislodgment rates differed between islands and mainland, controlling for colour type of the model (sample size = 3030 observations), and included “Insularity” (Island and Mainland habitat), “Model Dichromatism” (dichromatic vs. monochromatic) and their interaction as fixed effects. Here, “Model Dichromatism” captures how conspicuousness is distributed among individuals within a species: monochromatic species have similar male and female coloration, whereas dichromatic species have one sex, typically males, that is more conspicuous. This allows us to test whether species with sex-biased conspicuousness differ in overall predation risk.

For model 2, where we examined whether model colour contrast (green or brown contrast of the most conspicuous colour patch, based on V-type sensitivities) influenced dislodgment rates, we included as fixed effects “Insularity”, its interaction with “Chromatic contrast” and “Achromatic contrast”, and “NDVI” (vegetation density). Due to high correlations among different background contrast measures (green and brown r<0.85), to avoid collinearity issues (Freckleton 2010), we conducted two separate statistical models for each background colour (model 2a for brown contrast, model 2b for green contrast).

To explicitly test for sex-specific differences, i.e. being female or male model influenced the probability of being hit, we conducted a separate analysis restricted to dichromatic species (Model 3), which included “Insularity” and its interaction with “Sex-related Colour” (Female or Male), “Transect day” and “NDVI” as fixed effects (excluding all the monochromatic model species, sample size = 2074 observations).

Significant interaction terms were further examined using post hoc tests implemented in the emmeans package in R (Lenth 2025), including Tukey-adjusted pairwise comparisons of estimated marginal means and analyses of estimated marginal trends. All analyses were run in R (version 4.3.0, R Core Team 2024) using the lme4 package (Bates, Mächler et al. 2015). Data used for these analyses are available in the Dryad Digital Repository.

## Results

### Do hit rates differ between islands and mainland?

A total of 3.04% of all models deployed were dislodged (92 dislodgements out of the 3030 models placed in total, after exclusions). Of these, 1.15% (35) were dislodged on islands and 1.88% (57) on the mainland.

We found that models placed on islands experiencing lower dislodgment rates compared to those on the mainland (-1.098 [95% CI -2.150 – -0.045]); Figure 4). We did not observe differences between dichromatic and monochromatic models (Figure 4a; Supplementary Table S4 - model 1), but we found a significant interaction between insularity and model dichromatism on hit rate (2.038 [95% CI 0.198 – 3.879]); Figure 4b), indicating that the effect of insularity differed between model types. Post-hoc analysis indicated that dichromatic models experienced more hits on the mainland compared to the islands (z = 2.045, p = 0.04), while monochromatic models showed no significant difference between island and mainland, although a non-significant trend towards being more dislodged on islands compared to mainland (z = -0.941, p = 0.220, Figure 4b). Transect day was significant, with more dislodgments occurring on the second day of sampling (0.432 [95% CI 0.019 – 0.845])).

**Figure 4.**
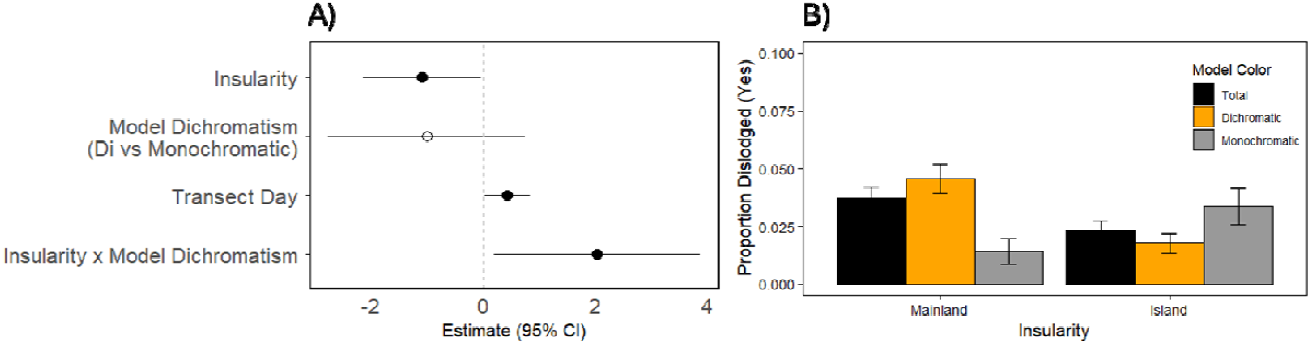
(**A**) Predictors of Dislodgment. The forest plot depicts effects (±95% CI) of each predictor on Dislodgment rates considering Insularity and Model Colour. Insularity is a categorical variable with Mainland set as the reference. Transect Day reference is Day 1. Model Colour reference is Dichromatic colour. Positive effects represent increases in dislodgment with increases of the explanatory variable. (**B)** Proportion of model dislodgments between island and mainland environments for the overall, dichromatic and monochromatic model colour type.

### Does conspicuousness relate with number of hits?

When analysing colour contrast against brown colours, we found a significant and positive effect of the interaction between insularity and chromatic contrast (-0.778 [95% CI -1.548 – - 0.009]); Figure 5). Post-hoc analysis showed that chromatic contrast tended to increase dislodgement probability on the mainland (slope = 0.400, p = 0.063) but showed no significant effect on islands (slope = -0.379, p = 0.276). The significant difference between these slopes (p = 0.047) confirms that chromatic contrast influences hit rates differently across environments (Figure 6). Transect day also had a significant effect, with Day 2 showing higher dislodgement rates compared to Day 1 (0.434 [95% CI 0.021 – 0.847]); Figure 5). All other factors were non-significant (Figure 5; Supplementary Table S4 - model 2a).

**Figure 5.**
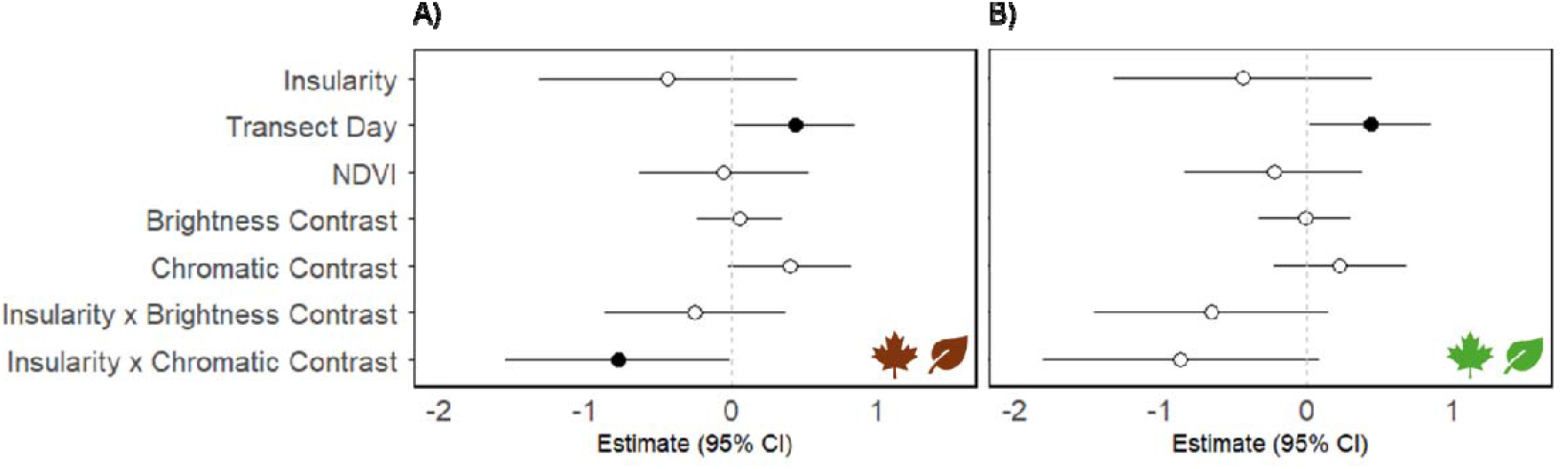
Predictors of Dislodgment. Forest plots depict effects (±95% CI) of each predictor on Dislodgment for analyses considering Brown (A) and Green background (B). Insularity is a categorical variable with Mainland set as the reference. Transect Day reference is Day 1. Positive effects represent increases in dislodgment with increases of the explanatory variable.

**Figure 6.**
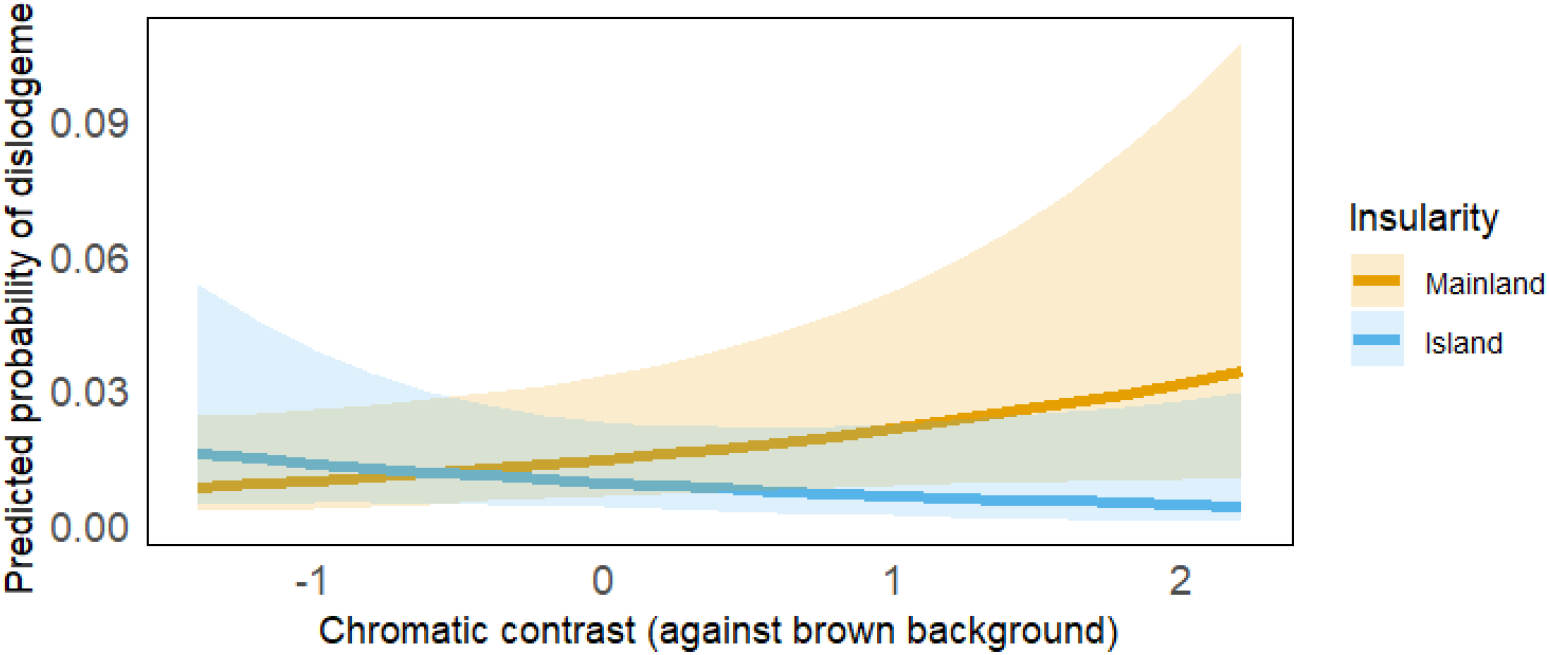
Predicted probability of dislodgement as a function of chromatic contrast against brown background. Lines show the predicted probability of a model being dislodged by predators as chromatic contrast (scaled) increases, with shaded ribbons representing 95% confidence intervals. Colours indicate insularity: blue for islands and orange for mainland sites. Predicted values of the interaction effect were derived using the ggeffects package in R (Lüdecke 2018).

When we analysed green background contrast, only Transect Day had a significant and positive effect but there was a tendency for a positive effect of the interaction between insularity and chromatic contrast (Figure 5, Supplementary Table S4 - model 2b).

### Are there more hits for males than females in dichromatic species?

For dichromatic model species results showed that the sex-related colour of the model (female or male model) did not influence the probability of being dislodged (alone and in interaction with insularity; Figure 7, Supplementary Table S4 - model 3). Vegetation and Transect day had also not effect (Supplementary Table S4 - model 3). In this model we found again a negative insularity effect, indicating that models placed on island had lower dislodgment rates compared to the mainland (-1.760 [95% CI -3.170–-0.351]); Figure 7).

**Figure 7.**
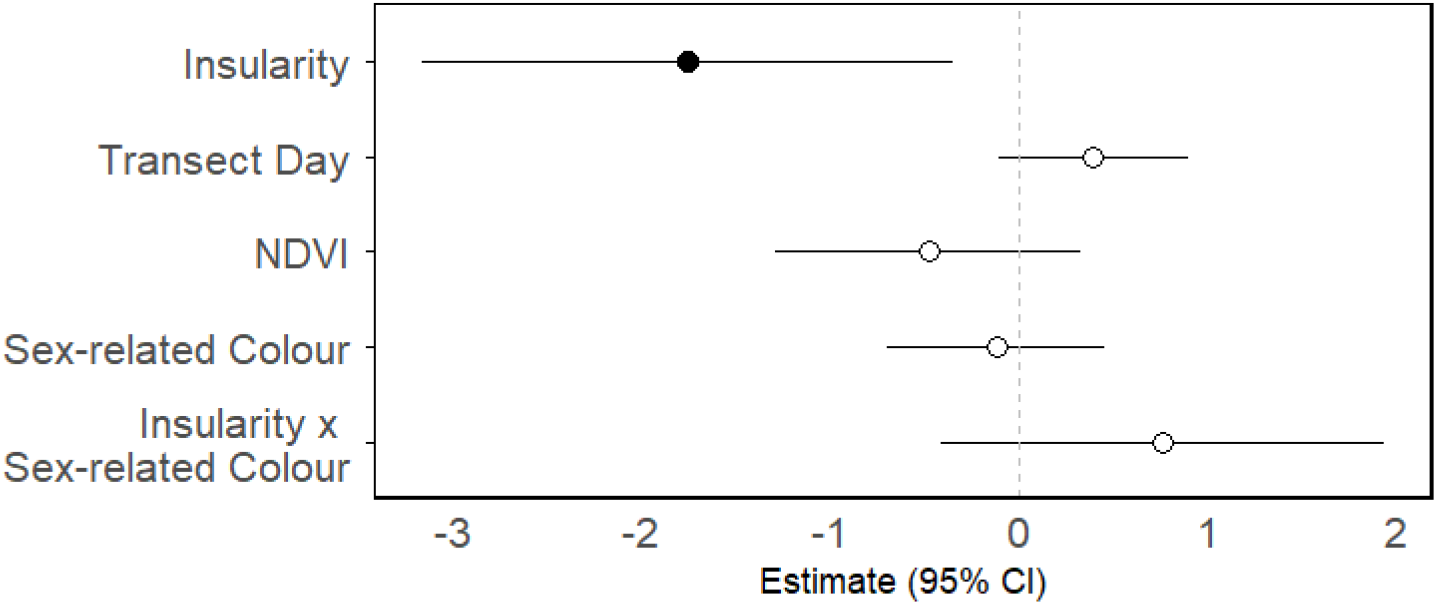
Predictors of Dislodgment. The forest plot depicts effects (±95% CI) of each predictor on Dislodgment analyses considering Sex-related colour, i.e. being female or male model, only for dichromatic species. Insularity is a categorical variable with Mainland set as the reference. Sex-related colour is categorical with Female set as the reference. Positive effects represent increases in dislodgment with increases of the explanatory variable.

### Putative predators and other detected animals by camera traps

A total of 12.5% of the camera traps (8 out of 64) captured a wild animal actively dislodging the model, including carnivores (*Vulpes vulpes, Herpestes ichneumon*), and suids (*Sus scrofa*). Birds of prey were not detected in our cameras, although in five cases, the cameras were set on models that were hit, but the animal responsible was not recorded. Of the 121 camera traps deployed, 52.9% (64 cameras) recorded wildlife passing by the model but not interfering with them, although they could have potentially been able to dislodge the models. These animals were birds (*Ardea cinerea, Turdus merula, Garrulus glandarius, Scolopax rusticola, Fringilla c. canariensis, Pyrrhula murina, Alectoris rufa*), squamate reptiles (*Gallotia atlantica, Tarentola angustimentalis*), rodents (possibly *Mus musculus, Rattus rattus* and/or *Rattus norvegicus)*, domestic dogs and cats, ungulates (*Capreolus capreolus*), bovids (*Capra hircus*), and other mammals (*Erinaceus europaeus, Atlantoxerus getulus*). In one notable case, a model was recognised by a conspecific, which responded with an aggressive display. Visual documentation is provided in Supplementary Figure S4.

## Discussion

In this study, we designed an experiment to test whether predation pressure differs between islands and the mainland and whether this variation is associated with differences in bird colouration, particularly focusing on conspicuousness and sexual dichromatism. Using 3D bird models, we assessed potential predation pressure through dislodgement rates. We found that models were hit less frequently on islands compared to the mainland, suggesting that islands have lower predation pressure. On the mainland, models with sexually dichromatic colouration and with higher chromatic contrast were more likely to be dislodged, indicating that more conspicuous models have increased predation risk.

We detected the predicted insularity effect, with significantly fewer dislodgements on islands compared to the mainland. This provides experimental support for the long-standing hypothesis that predation pressure is reduced in insular ecosystems. While predation levels may vary among islands depending on the type of predators present and densities, our results are consistent with expectations of reduced predation in insular environments, where predator richness and abundance are generally lower (Beauchamp 2004, Bliard, Paquet et al. 2020). This finding is also consistent with previous studies reporting relaxed anti-predator behaviours in island species (Blumstein and Daniel 2005, Cooper Jr, Pyron et al. 2014).

We also found that sexually dichromatic models, considering both female and male models, were more frequently dislodged on the mainland than monochromatic models, aligning with comparative studies showing increased predation risk in sexually dichromatic species (Huhta, Rytkönen et al. 2003, Møller and Nielsen 2006). Higher chromatic contrast was also positively associated with hit rates on the mainland, reinforcing the prediction that predators preferentially target more visible individuals (Götmark 1992, Vignieri, Larson et al. 2010, Troscianko, Wilson-Aggarwal et al. 2016). These findings suggest that sexual dichromatism and conspicuous colouration can carry survival costs under high predation pressure contexts, in line with theories of predator-driven selection against exaggerated signals (Endler 1978, Andersson 1994).

When examining sex differences in the models’ probability of being hit, we found no additional effects of sex-specific colouration beyond general colour contrast or dichromatism. This suggests that overall colour conspicuousness, rather than sexual signalling per se, is the main driver of predation risk in our system. One likely explanation is that sexual dichromatism in several of the species studied is relatively modest (Supplementary Figure S3), making male and female differences too small to influence predator decisions once overall contrast is accounted for. In addition, our models provided only static visual cues and predators may target males during active display behaviours that amplify ornamental traits, which were absent here. It is also possible that local predators respond mainly to general chromatic cues rather than fine-scale sex-specific patterns, or that the predator community may have limited sensitivity to subtle male and female differences. In all, these observations imply that the expression of sexual dimorphism may be strongly modulated by ecological and predation context: on islands with reduced predator pressure, relaxed selection may allow for reduced male and female differences, whereas on the mainland, sexual selection may interact with predation to maintain dimorphism despite its costs. Together, these factors suggest that male-biased predation may be contingent on behavioural and ecological factors not captured by our standardised colour models.

One constraint of our study is that overall dislodgement rates (3%), which reduces statistical power to detect subtle effects (such as sex-specific effects), although the main patterns remain robust. Such low rates are consistent with other artificial prey studies, which typically report hit rates between 3% and 18% across taxa and different settings (Supplementary Table S5). Such rates likely reflect the naturally low number of encounters between predators and models in the field, and the relatively short exposure time of these models. In addition, these artificial models lacked motion, behavioural displays, or acoustic cues, which are important components of sexual cues used by predators in natural conditions. Incorporating such cues in future studies could provide a more complete understanding of how sexual signals interact with predation risk. Nonetheless, the controlled approach used here also offers advantages. By holding behaviour constant and systematically varying specific traits and environmental contexts, we were able to isolate the effect of colouration on predation in a way that is often difficult to achieve in observational studies. While factors like ambient light, habitat complexity, and local predator communities may also influence predation patterns and warrant further investigation, our experimental design provides a complementary tool for disentangling the selective forces shaping visual signals in nature (see other similar studies on Supplementary Table S5).

Camera traps did not capture avian predation events, which was not entirely unexpected. Avian predation, particularly by birds of prey, are often underrepresented in camera-trap studies, as attacks may occur too quickly, predators may avoid cameras, or prey may be detected outside the camera’s view (Akcali, Pérez-Mendoza et al. 2019). Previous research shows that artificial prey experiments often detect bird predation through model damage rather than direct video confirmation (Akcali, Pérez-Mendoza et al. 2019, Schillé, Plat et al. 2025), and many studies of predation risk similarly rely on indirect indicators such as predator presence, or signs of attack rather than direct predation encounters (Supplementary Table S5). Our experimental approach, based on dislodgement of models as a proxy of predation, provides a standardized and comparable measure of predation risk, even if it cannot fully capture the identity or behaviour of the attacking species. Note that some dislodgements may have resulted from non-predatory disturbances, including rabbits or livestock. Although we excluded transects and models clearly affected by such events, the possibility of occasional undetected disturbance cannot be fully dismissed. Our data did not reveal consistent differences in the abundance of these disturbances between islands and the mainland. These events would likely introduce noise to the data rather than generate directional patterns, and would therefore be expected to weaken, rather than artificially create the differences we observed between islands and the mainland or between models of varying conspicuousness. Nonetheless, this potential source of uncertainty means that our estimates should be interpreted as reflecting relative differences in disturbance or predation risk under natural field conditions, rather than precise measures of predator-driven attacks alone. Additionally, while insular systems typically host fewer native predators (MacArthur and Wilson 1967, Beauchamp 2004), these systems often host a large numbers of introduced species (Bellard, Rysman et al. 2017). Separating the effects of native predators, introduced predators, or incidental disturbance was beyond the scope of this experimental design, yet our results still reflect how detectable models are under realistic ecological conditions, providing insights into how conspicuous traits may influence predation risk across environments.

Our findings suggest that predation is lower on islands and higher on the mainland, and that, across environments, increased conspicuousness is associated with higher predation risk. This supports the expected effect of conspicuousness increasing predation risk, but it also raises an apparent paradox: if predation is indeed relaxed on islands, why has this not favoured the evolution of more conspicuous colourations through other selective pressures? While some insular species do become more colourful under very low predation (Bliard, Paquet et al. 2020), other factors may constrain ornamentation. Reduced sexual selection, especially in small, isolated populations, may limit investment in costly traits, which can increase extinction risk (Doherty Jr, Sorci et al. 2003, Morrow and Pitcher 2003). Limited genetic variation (Griffith 2000, Whittaker and Fernández-Palacios 2007), shift toward K life history trait (Covas 2012) and energetic constraints in resource-limited environments such as islands may also favour the evolution of less conspicuous traits to signal quality. Note that our study does not test the evolutionary causes of these colour differences directly but rather evaluates whether predation pressure differs between environments and whether conspicuousness incurs differential risk. Overall, these findings suggest that relaxed predation alone is insufficient to create the ecological conditions favouring the evolution of conspicuous signals on islands, and that ecological, genetic, and life-history constraints are likely to modulate how insular populations diversify their visual traits.

In summary, our study provides empirical support that predation risk varies between islands and the mainland, and while predation clearly constrains conspicuousness where predator pressure is high, our results suggest that predation is not the main driver of the evolution of reduced colouration in island birds. This highlights the value of experimental island-mainland comparisons for testing long-standing evolutionary hypotheses and suggests that insular environments may serve as arenas for relaxed predation and increased signal diversification.

## Supplementary material

Supplementary material is available.

## Data and code availability

The data and code underlying this article are available in Dryad Digital Repository, at https://zenodo.org/records/18434436

## Author contributions

**Ana V. Leitão** (Conceptualization [lead], Methodology [lead]; Investigation [lead]; Resources [lead]; Data curation [lead]; Formal analysis [lead]; Funding acquisition [lead]; Writing - original draft [lead]; Writing - review and editing [lead]). **Cristina D. Alonso Moya** (Data curation [equal]; Investigation [equal]; Formal analysis [equal]; Writing - original draft [equal]; Writing - review and editing [supporting]). **Ricardo J. Lopes (**Resources [equal]; Methodology [supporting]; Writing - review and editing [supporting]). **Raquel Ponti ((**Resources [equal]; Data curation [supporting], Formal analysis [supporting], Writing - original draft [supporting]; Writing - review and editing [supporting]). **Rita Covas** (Conceptualization [equal]; Methodology [equal]; Resources [equal]; Formal analysis [supporting]; Funding acquisition [supporting]; Writing - original draft [supporting]; Writing - review and editing [supporting]). **Claire Doutrelant** (Conceptualization [equal]; Methodology [equal]; Resources [equal]; Formal analysis [supporting]; Funding acquisition [supporting]; Writing - original draft [supporting]; Writing - review and editing [supporting]).

## Acknowledgements

We are grateful to Juan Carlos Illera for his valuable assistance preparing fieldwork in the Canary Islands. We also thank David Gonçalves, Tiago Rodrigues, Luke Powell, Francisco Bonadonna and the PLT at CEFE for lending some of the camera traps. We thank Pedro Gonçalves from 3D Filamentos for all the support with the 3D printing, Osvaldo Branquinho for assistance with the figures, and the Behavioural Ecology and Sociality group members for valuable discussions. We are also grateful to all private landowners for allowing access to their land. This study was approved by the CIBIO’s Animal Welfare and Ethics Review Body (ORBEA). Fieldwork was conducted under licences from DRAAC of the Azores (2022/6800 & 2023/2346 - 005.01.02/31), Cabildo de Fuerteventura (2023/4072), Cabildo Insular de Tenerife (2023-00412), IFCN Madeira (5/2023M); permission for mainland Portugal was granted by ICNF without a licence requirement.

