## Supplementary material for "Predation and the Evolution of Island Bird Plumage Colouration: Experimental Insights from Island and Mainland Environments": Figure S1, Figure S2, Figure S3, Figure S4, Table S1, Table S2, Table S3, Table S4, Table S5

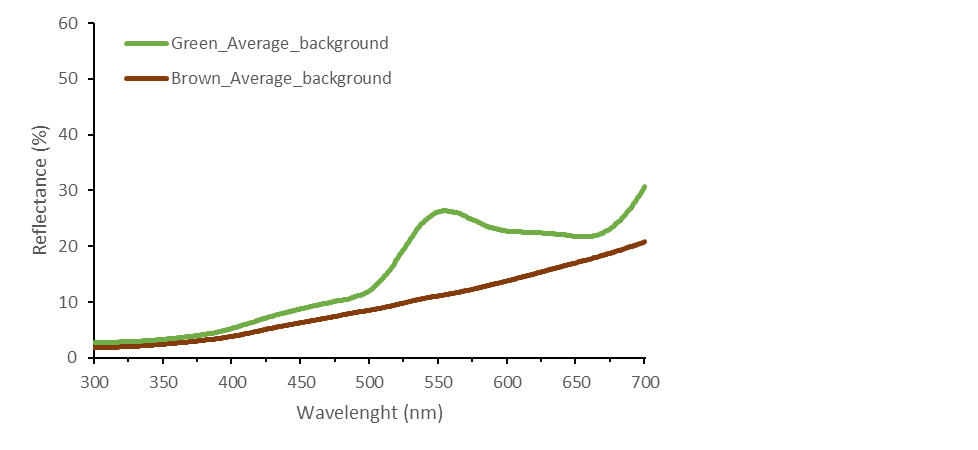

#### **Figure S1.** Reflectance spectra of the average background measurements. Lines show the average reflectance (%) of the two background categories measured in the field: green (leaves, grass, bushes) and brown (leaves, soil, bark). The x-axis represents wavelength (300–700 nm) and the y-axis represents reflectance percentage. These spectra were used to characterise the visual background for modelling chromatic and achromatic contrasts of the models.

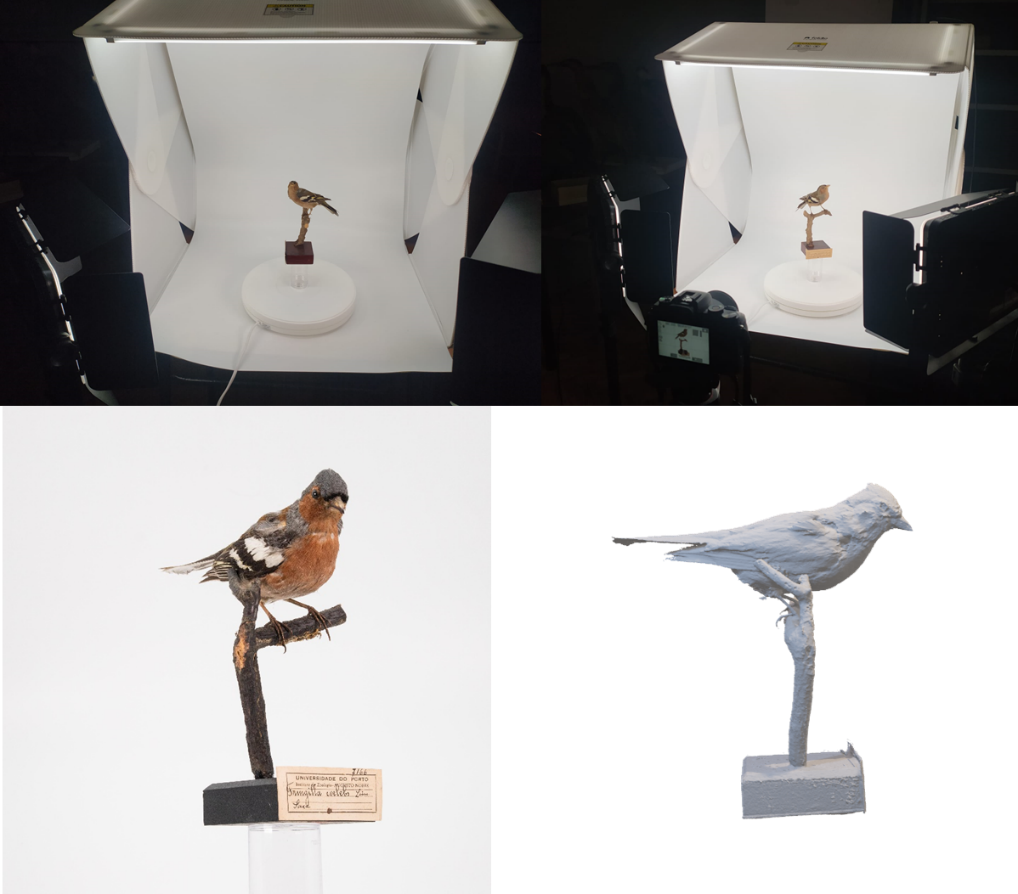

#### **Figure S2.** **Top photos**: Physical Setup for 360° Image Capturing. The specimen was positioned on a turntable, supported by a transparent stand to minimize visual obstructions. Lighting was supplied by two external spotlights, complemented by an integrated LED strip within the foldable chamber. Images were captured using a camera controlled remotely in continuous shooting mode to ensure consistency. **Bottom photos**: photo and converted 3D model file of the specimen. The specimens featured in this setup are a *Fringilla coelebs*.

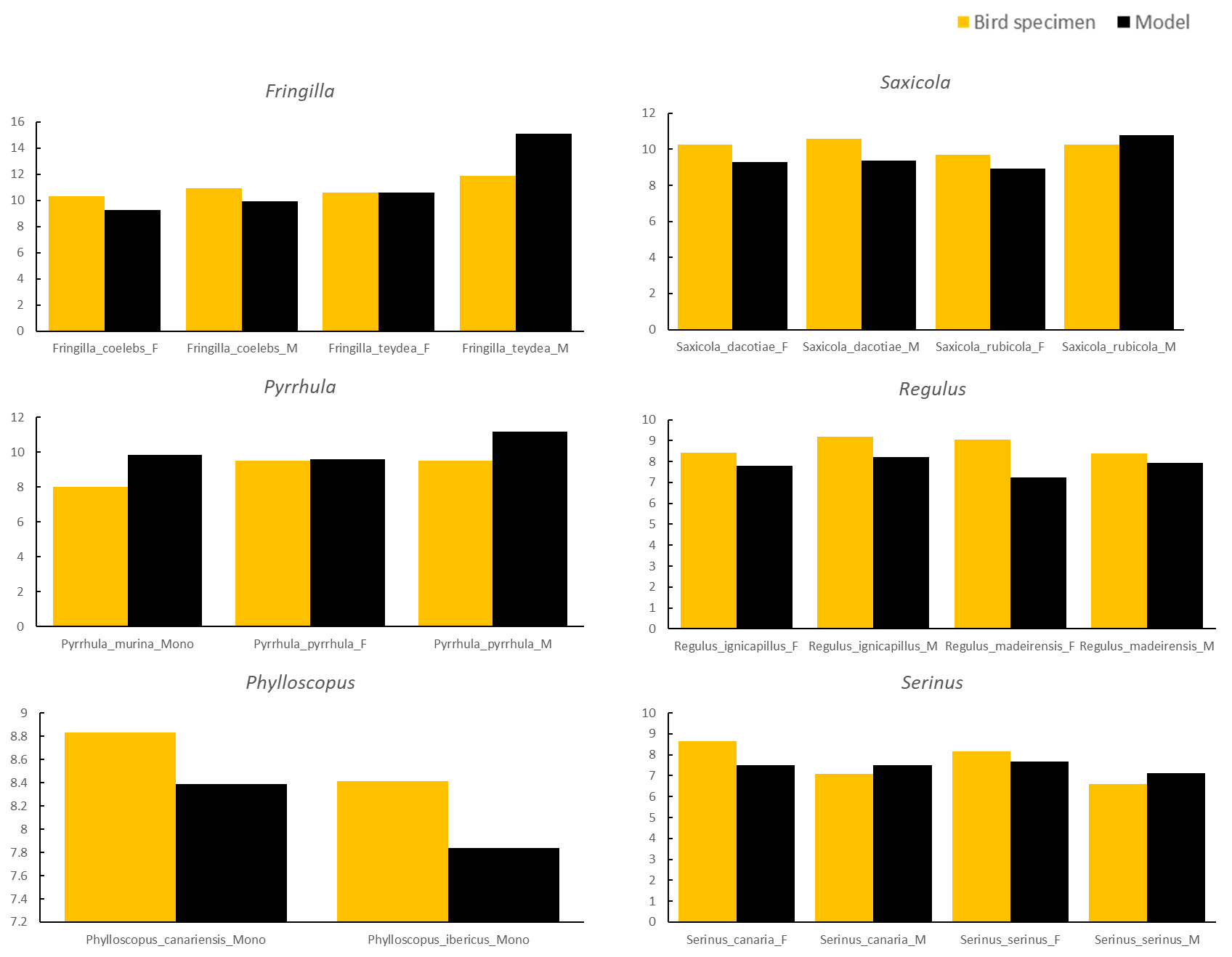

#### **Figure S3.** Chromatic contrast (∆S) of bird specimen plumage and painted models against natural background measured in receptor quantum catches.

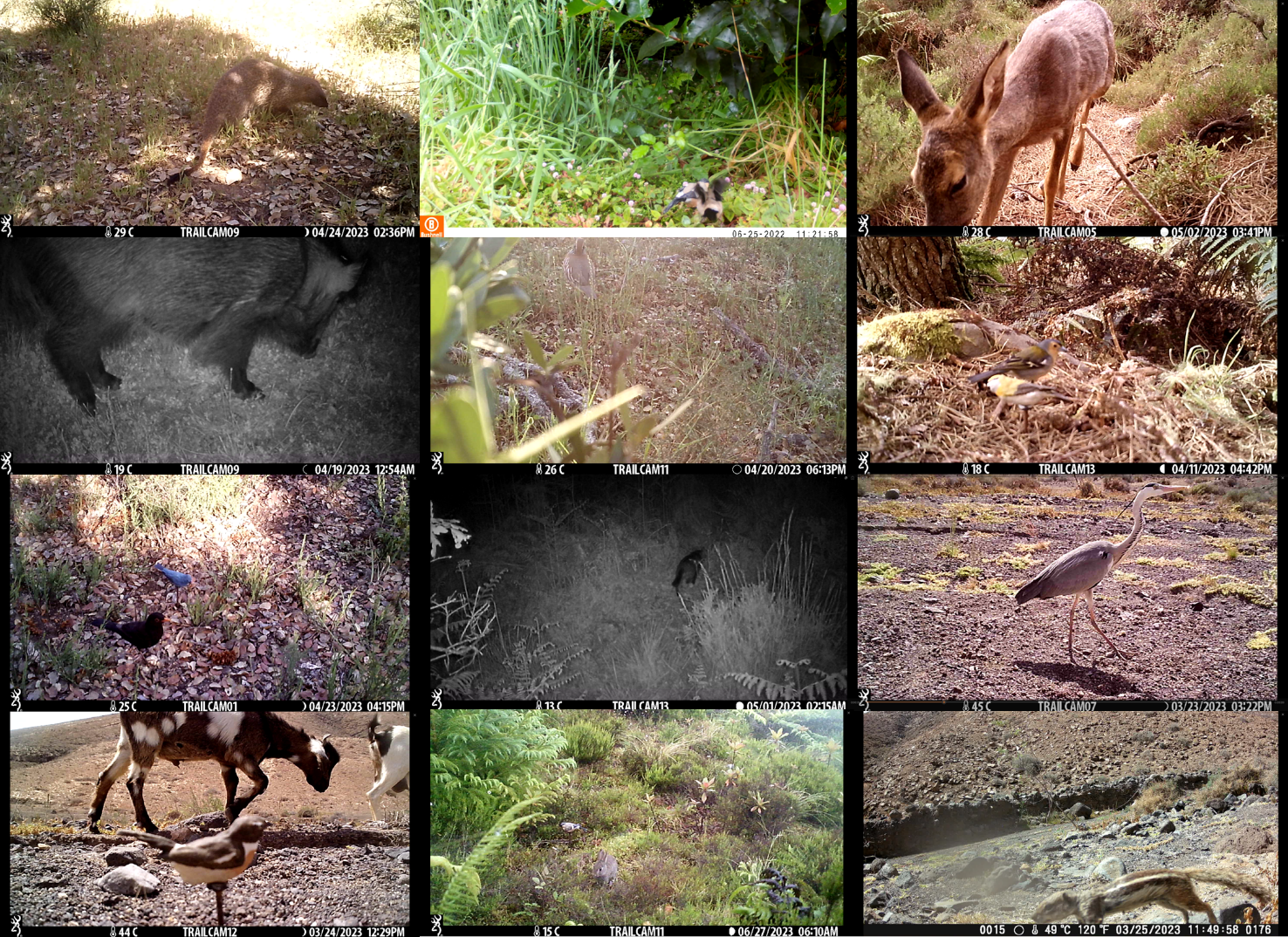

#### **Figure S4.** Wildlife caught on camera (not necessarily dislodging the models). Top to bottom, left to right: *Herpestes ichneumon*, *Pyrrhula murina, Capreolus capreolus, Sus scrofa, Alectoris rufa, Fringilla madeirensis, Turdus merula, Felis silvestris catus, Ardea cinerea, Capra hircus, Oryctolagus cuniculus, Atlantoxerus getulus*. Dates presented are month-day-year.

#### **Table S1.** Target species and information. Colours represent the different locations where the experiment was conducted: Green – Madeira; Yellow – Canary Islands; Blue – Azores.

| **Pair** | **IM** | **Common Name** | **Species Name** | **Family** | **Distribution** | **Field work location** | **♀♂** | **Breeding** | **Foraging location** |
| --- | --- | --- | --- | --- | --- | --- | --- | --- | --- |
| 1 | I | Madeira firecrest | *Regulus madeirensis* | Regulidae | Madeira, Portugal | Chão do Pasto & Ribeiro Frio | Dimorphic | April – July | Low to High height |
| 1 | M | Common firecrest | *Regulus ignicapilla* | Regulidae | Europe, N. Africa, and parts of Asia Minor | PNGerês, Portugal | Dimorphic | April – Aug | Low to High |
| 2 | I | Tenerife Blue Chaffinch | *Fringilla teydea* | Fringillidae | Canary Islands (Tenerife), Spain | Las Lagunetas, Santa Cruz de Tenerife | Dimorphic | April –August | Ground |
| 2 | M | Common chaffinch | *Fringilla coelebs* | Fringillidae | Western Palearctic and Central Russia | Portalegre, Alentejo, Portugal | Dimorphic | April –June | Ground |
| 3 | I | Canary Islands Chiffchaff | *Phylloscopus canariensis* | Phylloscopidae | Canary Islands (Tenerife), Spain | Las Lagunetas, Santa Cruz de Tenerife | Monom. | January - June | All levels |
| 3 | M | Iberian Chiffchaff | *Phylloscopus ibericus* | Phylloscopidae | Iberian Peninsula and N. Morocco | PNGerês, Portugal | Monom. | April - ? | All levels |
| 4 | I | Fuertventura stonechat | *Saxicola dacotiae* | Muscicapidae | Canary Islands (Fuerteventura) | Parque Natural Jandía, Fuerteventura | Dimorphic | Dec – April | Low to height |
| 4 | M | European stonechat | *Saxicola rubicola* | Muscicapidae | W&S Europe, N. Africa, Middle East | Portalegre, Alentejo, Portugal | Dimorphic | April – Aug | Low to height |
| 5 | I | Atlantic Canary | *Serinus canaria* | Fringillidae | Macaronesian islands | Chão do Pasto & Ribeiro Frio | Dimorphic | Canary Jan - July; Madeira Mar – Jun | Ground |
| 5 | M | European serin | *Serinus serinus* | Fringillidae | Europe, Middle East, N. Africa | Portalegre, Alentejo, Portugal | Dimorphic | April –July | Ground |
| 6 | I | Azores bullfinch | *Pyrrhula murina* | Fringillidae | Azores (São Miguel), Portugal | Pico Bartolomeu & Tronqueira | Monom. | June – August | Low - Medium |
| 6 | M | Eurasian bullfinch | *Pyrrhula pyrrhula* | Fringillidae | Europe and part of Asia | PNGerês, Portugal | Dimorphic | April – Aug | Low - Medium |

#### **Table S2.** Transects conducted at each site. Four transects were excluded from the analysis - two for *Serinus* and two for *Saxicola*, all on the island. The numbers shown do not include these excluded transects.

| **Genus** | **Island species** | **Island Site** | **Total nr transects days (24h)** | **Mainland species** | **Mainland Site** | **Total nr transects days (24h)** |
| --- | --- | --- | --- | --- | --- | --- |
| *Saxicola* | Fuerteventura stonechat | Canary Islands: Fuerteventura | 6 | European stonechat | Portalegre | 8 |
| *Pyrrhula* | Azores bullfinch | Azores: São Miguel | 6 | Eurasian bullfinch | PNGerês | 6 |
| *Serinus* | Atlantic Canary | Madeira | 4 | European serin | Portalegre | 6 |
| *Phylloscopus* | Canary Islands Chiffchaff | Canary Islands: Tenerife | 8 | Iberian chiffchaff | PNGerês | 6 |
| *Fringilla* | Blue Chaffinch | Canary Islands: Tenerife | 8 | Common chaffinch | Portalegre | 8 |
| *Regulus* | Madeira firecrest | Madeira | 6 | Common firecrest | Portalegre | 6 |
| Total transect days |  |  | 38 |  |  | 40 |

#### **Table S3.** Maximum chromatic and achromatic contrasts against brown and green for each model species. Colour contrasts are expressed in units of JNDs, where 1 JND is the theoretical threshold of colour discrimination. These distances indicate to what extent the different patches resemble natural backgrounds in colour, with higher values indicating more contrasting colours.

| **Model species** | **Habitat** | **Sex** | **Brown background** | | **Green background** | |
| --- | --- | --- | --- | --- | --- | --- |
|  |  |  | **Chromatic contrast** | **Achromatic contrast** | **Chromatic contrast** | **Achromatic contrast** |
| *Fringilla coelebs* | Mainland | F | 5.200 | 20.445 | 11.039 | 22.656 |
|  |  | M | 9.145 | 19.680 | 14.173 | 21.891 |
| *Fringilla teydea* | Island | F | 5.255 | 13.712 | 10.828 | 15.923 |
|  |  | M | 10.598 | 8.2512 | 15.463 | 6.886 |
| *Phylloscopus ibericus* | Mainland | Monomorphic | 3.355 | 17.998 | 8.855 | 20.209 |
| *Phylloscopus canariensis* | Island | Monomorphic | 3.398 | 19.186 | 8.954 | 21.398 |
| *Pyrrhula pyrrula* | Mainland | F | 6.955 | 15.598 | 11.283 | 17.810 |
|  |  | M | 6.588 | 15.751 | 12.014 | 17.962 |
| *Pyrrhula murina* | Island | Monomorphic | 6.857 | 16.837 | 12.333 | 14.625 |
| *Regulus ignicapilla* | Mainland | F | 7.619 | 18.351 | 11.228 | 20.563 |
|  |  | M | 9.743 | 14.952 | 11.173 | 17.163 |
| *Regulus madeirensis* | Island | F | 7.641 | 17.155 | 11.344 | 19.366 |
|  |  | M | 7.437 | 17.526 | 11.112 | 19.738 |
| *Saxicola rubicola* | Mainland | F | 5.435 | 19.593 | 10.893 | 21.804 |
|  |  | M | 6.001 | 17.843 | 11.339 | 20.055 |
| *Saxicola dacotiae* | Island | F | 5.986 | 18.919 | 11.315 | 21.130 |
|  |  | M | 5.274 | 21.143 | 10.754 | 23.354 |
| *Serinus serinus* | Mainland | F | 3.789 | 17.916 | 9.8833 | 20.127 |
|  |  | M | 7.273 | 17.147 | 9.7641 | 19.358 |
| *Serinus canaria* | Island | F | 3.988 | 13.158 | 9.8199 | 15.369 |
|  |  | M | 5.1716 | 14.084 | 9.9047 | 16.295 |

#### **Table S4.** Generalized mixed models assessing the relationship between the Dislodgment (yes/no) as the response variable with a binomial family and included as a fixed effect Insularity, Transect day, and different colour predictors.

|  | **Predictor** | **Estimate** | **SE** | **z** | **P** | **95% CI (lower; upper)** |
| --- | --- | --- | --- | --- | --- | --- |
| **Model 1** | Insularity (Mainland vs Island) | -1.098 | 0.536 | -2.045 | **0.048** | **(-2.150 ; -0.045)** |
|  | Model Dichromatism (Mono. vs Dichro.) | -1.008 | 0.895 | -1.126 | 0.260 | (-2.764 ; 0.747) |
|  | Transect Day (1^st^ vs 2^nd^ day) | 0.432 | 0.210 | 2.051 | **0.043** | **(0.019 ; 0.845)** |
|  | Insularity * Model type | 2.039 | 0.938 | 2.172 | **0.029** | **(0.198 ; 3.879)** |
|  |  | Variance | STD |  |  |  |
|  | Transect number (N=39) | 1.013 | 1.006 |  |  |  |
|  | Pair (N= 6) | 0.434 | 0.659 |  |  |  |
| **Model 2a** | Insularity (Mainland vs Island) | -0.437 | 0.449 | -0.972 | 0.331 | (-1.319 ; 0.444) |
|  | NDVI | -0.053 | 0.297 | -0.180 | 0.856 | (-0.637 ; 0.530 |
|  | Brightness contrast brown | 0.054 | 0.146 | 0.371 | 0.710 | (-0.233 ; 0.342 |
|  | Chromatic contrast brown | 0.399 | 0.215 | 1.854 | 0.063 | (-0.022 ; 0.822) |
|  | Transect Day (1^st^ vs 2^nd^ day) | 0.434 | 0.210 | 2.062 | **0.039** | **(0.021 ; 0.847)** |
|  | Insularity * Brightness | -0.252 | 0.316 | -0.797 | 0.425 | (-0.872 ; 0.367) |
|  | Insularity * Chromatic | -0.778 | 0.392 | -1.984 | **0.047** | **(-1.548 ; -0.009)** |
|  |  | Variance | STD |  |  |  |
|  | Transect number (N=39) | 1.108 | 1.052 |  |  |  |
|  | Pair (N= 6) | 0.460 | 0.678 |  |  |  |
| **Model 2b** | Insularity (Mainland vs Island) | -0.440 | 0.453 | -0.971 | 0.331 | (-1.329 ; 0.448) |
|  | NDVI | -0.226 | 0.311 | -0.727 | 0.467 | (-0.836 ; 0.383) |
|  | Brightness contrast green | -0.014 | 0.159 | -0.089 | 0.929 | (-0.326 ; 0.298) |
|  | Chromatic contrast green | 0.225 | 0.230 | 0.980 | 0.327 | (-0.225 ; 0.677 ) |
|  | Transect day (1^st^ vs 2^nd^ day) | 0.434 | 0.210 | 2.061 | **0.039** | **(0.021 ; 0.847)** |
|  | Insularity * Brightness | -0.650 | 0.408 | -1.593 | 0.111 | (-1.451 ; 0.149) |
|  | Insularity * Chromatic | -0.866 | 0.482 | -1.795 | 0.072 | (-1.812 ; 0.079) |
|  |  | Variance | STD |  |  |  |
|  | Transect number (N=39) | 1.137 | 1.066 |  |  |  |
|  | Pair (N= 6) | 0.368 | 0.606 |  |  |  |
|  | **Predictor** | **Estimate** | **SE** | **z-value** | **P** | **95% CI (lower; upper)** |
| **Model 3** | Insularity (Mainland vs Island) | -1.760 | 0.719 | -2.448 | **0.014** | **(-3.170 ; -0.351)** |
|  | NDVI | -0.482 | 0.414 | -1.165 | 0.243 | (-1.294 ; 0.329) |
|  | Sex-related colour (Female vs Male) | -0.121 | 0.294 | -0.411 | 0.681 | (-0.698 ; 0.456) |
|  | Transect day (1^st^ vs 2^nd^ day) | 0.395 | 0.257 | 1.537 | 0.124 | (-0.108 ; 0.898) |
|  | Insularity * Model Sex-related colour | 0.761 | 0.598 | 1.272 | 0.203 | (-0.411 ; 1.933) |
|  |  | Variance | STD |  |  |  |
|  | Transect number (N=26) | 1.129 | 1.062 |  |  |  |
|  | Pair (N= 4) | 0.280 | 0.529 |  |  |  |

**Table S5.** Summary of some experimental predation studies across different taxa. Species or taxonomic group, material used for the models, hit rates, model types, and methods of predator confirmation reported in previous studies.

| **Species/group** | **Material** | **Hit Rate** | **Confirmation of predators** | **Study** |
| --- | --- | --- | --- | --- |
| Birds  Coraciiformes and Upupiformes | Plaster | 5.6% hits 111 out of 1988 (all raptors) | Direct: video camera | (Ruiz-Rodríguez, Avilés et al. 2013) |
| Birds  *Malurus spp* | 3D-printed | 13.2% hits (331 out of 2511 models) | Direct and indirect: camera traps and dislodgment. | (Cain, Hall et al. 2019) |
| Mice  *Peromyscus polionotus* | Plasticine | 3% hits (28/892) | Indirect: Presence of marks, or moved | (Vignieri, Larson et al. 2010) |
| Frogs  *Oophaga pumilio* | Clay | 10.5% hits (283/2700) | Indirect: marks | (Preißler and Pröhl 2017) |
| Turtles  *Terrapene carolina* | 3D-printed | 18% interactions (65/372 trials; 8% by predators) | Direct: camera traps | (Tetzlaff, Estrada et al. 2020) |
| Snake  *Crotalus lepidus l.* | Foam models | 13% disturbed (55/421; 27 avian, 28 non-predators) | Indirect: marks. | (Farallo and Forstner 2012). |
| Lizard  *Ctenophorus decresii* and *C. vadnappa* | Plaster | 5.1 % dislodgment (113/2200) | Indirect: movement or dislodgment | (Stuart-Fox, Moussalli et al. 2003) |
| Beetles  Family: Cerambycids | Plasticine and cover | 13% hits (177/1360) | Indirect: Beak marks | (Goßmann, Ambrožová et al. 2023) |
